# Sperm competition influences sperm quality and seminal plasma protein expression among territorial males of the courtship cichlid

**DOI:** 10.1101/2023.09.07.556763

**Authors:** Masaya Morita, Shun Satoh, Takeshi Ito, Masanori Kohda, Satoshi Awata

## Abstract

Sperm competition drives the plasticity of sperm quality and seminal fluid. However, how environmental and behavioral factors influence sperm competition and its outcome on the plasticity of sperm quality are unknown. In this study, we show males can asses the sperm competition level, and their fertilization-promoting sperm traits changes according to intensity. The promiscuous Tanganyikan cichlids *Ophthalmotilapia ventralis* perform their fertilization in the female mouth cavity. Sperm competition occurs in the female mouth as sperm is collected from multiple males. Behavioral (i.e., courtship success rates [CSR]) and environmental (density of spawning sites [DS]) factors are related to sperm collection. Seminal plasma glycoprotein 120 (SPP120) aids in sperm immobilization and aggregation, and sperm quality (longevity and velocity) could increase the probability of siring offspring. Based on an assumption of the involvement of DS and CSR on sperm collection, DS and CSR could influence the expression of SPP120 and sperm quality. We examined how sperm quality and SPP120 affect sperm competition-associated CSR and DS. Sperm longevity and velocity were positively correlated with DS. SPP120 expression was also positively correlated with CSR and DS. The plasticity of sperm quality and amount of SPP120 result in high fertilization competency according to sperm competition level.

## Introduction

Female multiple mating often influences the direction and intensity of sexual selection (Barbosa et al., 2010; Birkhead and Pizzari, 2002; Eberhard, 1996; Sugg and Chesser, 1994; Urbach et al., 2005; Ward, 2007). Among traits subjected to sexual selection, gametic involved in fertilization success are strongly influenced by selection. For example, sperm characteristics and seminal plasma proteins (SPP) from competing ejaculates could be candidates for strongly selected traits if sperm compete for a limited number of ova, i.e., sperm competition (Clark and Swanson, 2005; Fitzpatrick et al., 2009; Iwata et al., 2011; Parker, 1998; Ramm et al., 2009; Ramm et al., 2008; Sirot et al., 2015), and if females manipulate sperm for fertilization after mating, i.e., cryptic female choice (Birkhead and Pizzari, 2002; Firman et al., 2017).

Sperm quality and the amount of SPP are predicted to be influenced by the levels of sperm competition and cryptic female choice. Sperm quality is correlated with fertilizing ability when sperm competition arises (Bencic et al., 1999; Levitan, 2000), and non-chosen males need to have tactics to splash-disperse their sperm during mating with the chosen partner (Taborsky, 1998). Within species, sperm motility (representative of sperm quality) changes according to the level of sperm competition. For example, faster swimming sperm have empirically shown their competitive ability to fertilize in teleost fish (Gage et al., 2004; Gage et al., 1995; Leach and Montgomerie, 2000; Ota et al., 2010; Stoltz and Neff, 2006; Vladic and Jarvi, 2001). Moreover, changes in SPP levels assist the ejaculated sperm in the fertilization process, as reported in *Drosophila* (Fedorka et al., 2011; Misra and Wolfner, 2020; Sirot et al., 2011; Wigby et al., 2009) and mice (Ramm et al., 2015). Among species, sperm competition is strongly correlated with the evolution of sperm motility (Fitzpatrick et al., 2009; Locatello et al., 2007). Studies have shown that sperm morphology (Simmons et al., 1999) and SPP have evolved rapidly to increase fertilization success (Dorus et al., 2004; Ramm et al., 2009). Many studies have also indicated the significance of seminal fluid and sperm motility in increasing fertilization success in response to sperm competition and cryptic female choice; however, the degree to which environmental factors and male attractiveness can change sperm motility and the amount of SPP remains unknown.

Cryptic female choice and sperm competition often arise simultaneously during internal fertilization due to multiple mating called “female promiscuity.” In external fertilization, cryptic female choice is often ignored because fertilization occurs without the female’s control, although there are some exceptions; for example, ejaculated sperm motility may be controlled by ovarian fluid from the female (Alonzo et al., 2016; Gasparini and Pilastro, 2011; Johnson et al., 2020). However, cryptic female choice and sperm competition can also arise in external fertilization in fish, demonstrating a unique spawning manner (e.g., Fitzpatrick, 2020). In the mouthbrooding cichlids *Ophthalmotilapia ventralis* endemic to Lake Tanganyika, fertilization occurs in the enclosed mouth cavity of females that collect sperm from multiple males (sperm competition); females may choose fertilization from mixed sperm (cryptic female choice) (Fig.1a, Haesler et al., 2011). Although the mouth cavity is not an internal region and sperm can quickly diffuse (Fig. 1a, Morita et al., 2018) due to their motility initiated after ejaculation (Morisawa and Suzuki, 1980; Morita et al., 2014), the seminal plasma glycoprotein 120 (SPP120), a glycosylated SPP, facilitates sperm to remain quiescent in the mouth without initiating motility (Fig. 1a and b, Mochida et al., 1999; Mochida et al., 2002; Morita et al., 2018). The female collects sperm before, during, and after spawning eggs at a male’s nest, called a sandy patch (“sperm shopping”) (Haesler et al., 2009; Immler and Taborsky, 2009; Morita et al., 2018), and then sperm from many males are stocked in the mouth cavity (sperm competition, Fig. 1a). In this scenario, cryptic female choice is suspected since the fertilization success is highly skewed toward particular males, and even those not included in sperm shopping can successfully sire some eggs during fertilization in the female mouth (Haesler et al., 2011). Although the cryptic female choice is implied, sperm competition is modeled as a “fair raffle,” and the number of sired offspring depends on the relative sperm number in the “fertilization set” (Parker, 1990).

**Fig. 1:**
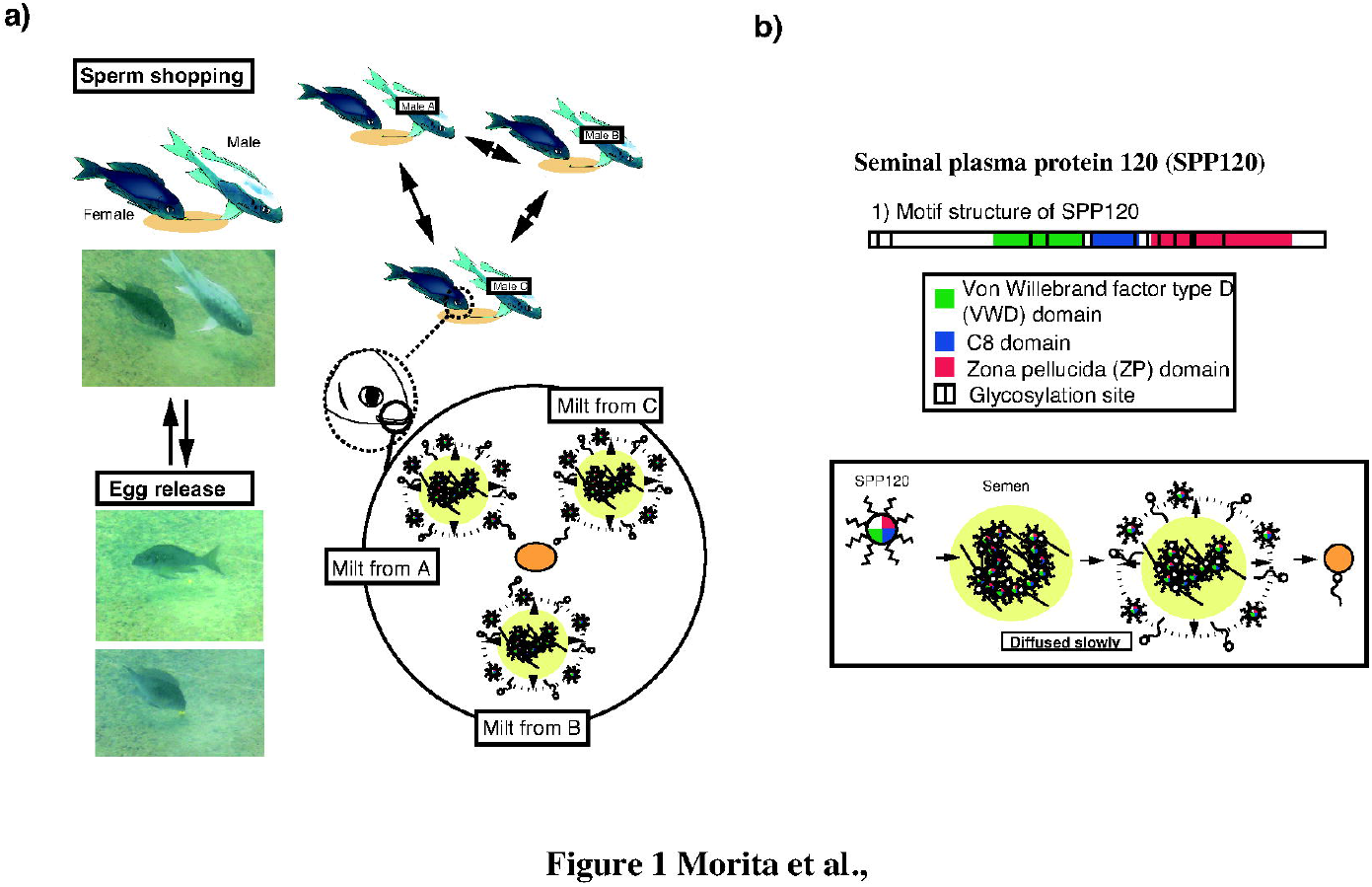
Spawning sequence and seminal plasma protein 120 (SPP120) role in the mouthbrooding cichlid *Ophthalmotilapia ventralis*. a) Sequence of courtship, sperm shopping, and egg release. The semen from many males is collected in the female mouth cavity, and fertilization occurs with sperm chosen from the stocked semen (high sperm competition level and cryptic female choice). b) SPP120 structure; three domains lead to sperm aggregation, and the N-terminal region immobilizes sperm flagellar motility. c) Predicted high-sperm-competition situation in the female mouth cavity after sperm shopping from multiple males and egg release.

Male attractiveness and patch density are implicated in sperm shopping and are with sperm competition (Fig. 1a). For example, more attractive males in denser patches cause higher sperm competition levels than those in less dense sandy patches. Furthermore, parasitic spawning by non-territorial males requires additional tactics for sperm competition. According to these behavioral traits and environmental conditions reflecting sperm competition, it is unknown whether sperm quality and SPP120 levels change to increase fertilization success. Nevertheless, the plasticity of sperm quality and amount of SPP120 expression could be a male tactic to oppose sperm competition and cryptic female choice.

To elucidate behavioral and environmental factors associated with sperm quality and SPP120 expression, we estimated male attractiveness as an index of courtship success rates (CSR) and risk of parasitic spawning as an encounter rate of non-territorial floating males (ERFM). The density of sandy patches (DS) was considered an environmental factor. For assessing sperm quality, sperm longevity, and swimming speed were measured, and SPP120 expression was evaluated quantitatively. It has been hypothesized that the risk of sperm competition increases with increasing DS and CSR in males or the number of floating males. Here, we examined the correlation between sperm quality and SPP120 expression and assessed the risk of sperm competition.

## Materials and Methods

### Animals

Thirty territorial males of *O. ventralis*, which had sandy patches for spawning, were used for observing behavioral traits, comparing testis size, observing sperm motility, and measuring SPP120 expression and amount. The sandy patches were localized at 1.9–3.8 m depth around Nkumbla Island in the southern part of Lake Tanganyika in the Republic of Zambia. Individual cichlids were identified based on black spots on the body or tagged with visible implant elastomer tags (Northwest Marine Technology, Inc., WA, USA). Animal experiments complied with the regulations of the Guide for Care and Use of Laboratory Animals. They were approved by the Committee of Laboratory Animal Experimentation at the University of the Ryukyus (21-01).

### Observation of reproductive behavior

Fish behavior was observed for 15 min/day over 10 days (total = 150 min/fish), with a focus on courtship success rates (CSR) and encounter rates of floating males (ERFM). We identified territorial males by black-spotted color or elastomer tags. CSR was measured as the number of female visits to a sandy patch in response to male courtship behavior. ERFM was categorized as positive if the male noticed and attacked the floating male, and then counted the number of the attacks during 15 min. All of the observed data were normalized. The encounter rate among territorial males was also observed, but territorial males maintained their territories without any overlap (e.g., Kohda, 1997) and thus rarely attacked other territorial males. Therefore, this category was excluded from the present study. However, the distance between the sandy patches of territorial males was measured.

### Different distances of the sandy patches

From spawning sequences, females collect sperm from territorial males multiple times, and distances and numbers of the patches could influence the sperm competition level. Therefore, we categorized the density of the patches as representative of the sperm competition level. We categorized the density of sandy patches (DS) as follows: n*{1/d_1_+ 1/d_2_+ 1/d_3_+ 1/d_n_} = n*∑1/d_n_, where d is the distance between sandy patches, and n is the number of sandy patches (Supplementary Fig. 1). Sandy patches located >10 m apart were excluded, as most females visited neighboring sandy patches within <10 m.

### Measurements of testes size and sperm motility

*O. ventralis* (*n* = 30) were captured at Nkumbla Island using a gill net after 10 days of observation. In the laboratory, the males were killed by an overdose of the anesthetic FA100 (Tanabe Seiyaku Inc., Osaka, Japan), weighed (to the nearest 0.001 g), and dissected, and their testes were weighed to the nearest 0.001 g. The mass of the testes was measured to calculate the gonadosomatic index (GSI: total testes mass/body mass × 100) to estimate the intensity of sperm competition and its correlation with sperm longevity. The testes were then dissected to collect sperm for observation of motility. Sperm motility was determined according to the method described by Morita et al. (2014). We then assessed sperm longevity and swimming speed 10 seconds after sperm motility was initiated via suspension in lake water.

### RNA isolation and cDNA synthesis

Total RNA was extracted from 1 cm^2^ tissue sections according to the method described by Morita et al. (2018). Testes from 24 individuals were first fixed using the RNA stabilization solution RNAlater (Ambion) and stored at -20 °C. Then, total RNA was isolated using TRIpure Reagent (Roche, Basel, Switzerland). The testes samples were homogenized with TRIpure reagent and centrifuged at 10,000 *g* for 10 min at four °C. The supernatant was mixed with 0.2 (v/v) chloroform, and centrifuged at 10,000 *g* for 10 min at 4 °C. The supernatant was separated, and then an equal volume of 2□propanol was added to the precipitate of total RNA; the sample was centrifuged again 10,000 *g* for 10 min at 4 °C. Total RNA pellets were washed with 75% ethanol, dried, and resuspended in 30 μL nuclease-free water. RNA was quantified spectrophotometrically using an ND□1000 UV/visible spectrophotometer (NanoDrop Technologies). The TURBO DNA□free™ Kit (Applied Biosystems, Foster City, CA, USA) was used to remove contaminating DNA from total RNA, and cDNA was synthesized using SuperScript III (Invitrogen, Carlsbad, CA, USA) with random hexamers.

### Isolation of the partial sequence of *EF1*α and the sequences of SPP120

A partial sequence of the translation elongation factor (EF1a) was isolated to normalize the gene expression level. Based on the complete sequences of *EF1*α obtained from teleost, a highly conserved sequence was selected to design primers for PCR. The forward primer sequence was 5’-CACA(TorA)(CorT)AACATCGTGGTCAT(TorC)-3,’ and the reverse primer sequence was 5’-GGGTGGTTCAGGATGATG-3.’ PCR was performed using cDNA templates and involved 35 cycles with the following steps: 92 °C for 30 s, 57 °C for 30 s, and 72 °C for 2 min for *EF1*α or 3 min for *SPP120*. The PCR products were separated on 1% agarose gel by electrophoresis. The PCR product was ligated, cloned into the pGEM-T Easy Vector (Promega, Madison, WI, USA), and sequenced.

### *SPP120* expression analysis

To analyze the expression level of *SPP120*, real-time PCR was performed in a reaction volume of 20 μL comprising 10 μL Mastermix with SYBR green (Bio-Rad, Hercules, CA, USA), 0.6 μL each primer (1.2 μM), 0.3 μL sample cDNA (2.5 µg total RNA/ 20 µL), and 8.5 μL ultrapure water. Reactions were run on an iCycler (Bio-Rad, Japan) under the following conditions: 1 cycle at 95 °C for 3 min; 45 cycles of 94 °C for 30 s, 55 °C for 30 s, and 72°C for 30 s; and 1 cycle of 95 °C for 15 s, 60 °C for 30 s, and 95 °C for 15 s. Data were normalized to the expression level of *EF1α*.

Primers were designed with Primer 3 (v. 0.4.0) software based on the complete sequences of *SPP120_*01 and *SPP120_*02 (functional genes), *SPP120_*03 and *SPP120_*04 (pseudogenes) (Supplementary Fig. 2), and the partial sequence of *EF1α*. We used *SPP120_*01 to analyze the expression of *SPP120, SPP120_*03, and *SPP120_*04 according to the criteria described in Supplementary Information 1.

### Statistical analysis

The correlations between behavioral and environmental parameters (ERFM, CSR, and DS), sperm quality (sperm longevity and swimming speed), and *SPP120* expression were analyzed using the linear model method via the lm functions of R, implemented in the “nlme” package. The normality and homogeneity of the data variance were checked using the Shapiro–Wilk and Bartlett tests, respectively, before analysis. If normality was rejected, the data were log-transformed. First, we analyzed a complete modeled set of gametic traits (*SPP120* expression, sperm longevity, and swimming speed) as an objective. We set ERFM, CSR, DS, and their three-way interaction as explanatory variables in a full model of separate linear models (LMs). Then, the likelihood ratio (LR) test was used to select explanatory variables to drop. Next, the updated formula was determined, and the significance of the difference between models before and after dropping was tested. Then, variables were dropped until no additional parameters to remove from the formula were identified. After the final models were determined, it was recalculated using the restricted maximum likelihood method. All analyses were performed using the R ver. 4.0.2.

## Results

### Polyandry and parasitic spawning

During the study, spawning was observed three times with durations of 952, 1710, and 4365 s. Females collected sperm from two to seven males before, during, and after spawning eggs (Supplementary Information 2). The time between sperm shopping and the spawning of eggs varied (Fig. 2a; Supplementary Information 2). Moreover, the number of eggs released was smaller than that of sperm (Fig. 2b). Females did not deposit eggs at sandy patches of several males, despite collecting sperm from these males multiple times (Supplementary Information 2). Parasitic spawning was also observed via the sneaking behavior of floating males, representing one-third of the observed spawning events (Fig. 2a, Supplementary Information 2).

**Fig. 2:**
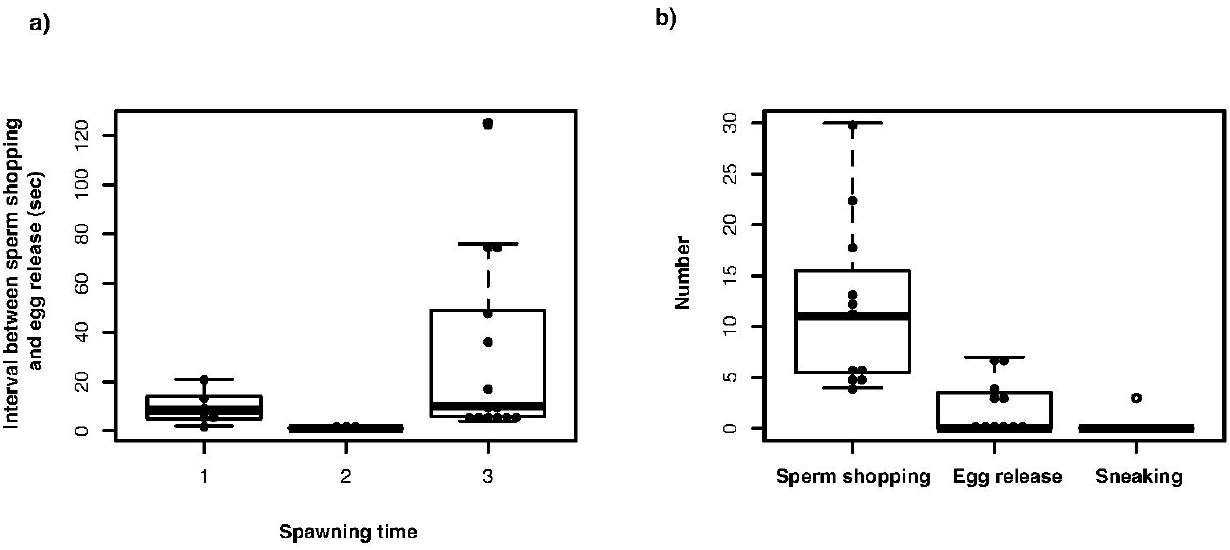
Observed spawning parameters in *Ophthalmotilapia ventralis*. a) Intervals of sperm shopping and egg release at each observed spawning. b) Frequencies of sperm shopping, egg release, and parasitic spawning (sneaking) by floating males.

### Floating males, the density of sandy patches, and courtship success rates

None of the behavioral and environmental traits (ERFM, DS, and CSR) were correlated (Supplementary Table 1).

### GSI, sperm traits, and *SPP120* expression

GSI and fertilization-related traits (sperm motility and *SPP120* expression) did not differ significantly between males with whom females spawned eggs and those with whom females did not spawn (Supplementary Fig. 3a, b, c, and d; Wilcoxon rank sum test). In contrast, DS, CSR, and ERFM were associated with fertilization-related traits and GSI (Table 1 and Fig. 3a).

**Table 1.** Correlation of gametic traits and GSI with behavioral parameters from linear model.

| Trait | Predictors | Coefficient | S.D. | t | p |
| --- | --- | --- | --- | --- | --- |
| **Sperm longevity** | **Floating male (ERFM)** | 0.19 | 0.06 | 3.1 | 0.005 |
|  | **Density of sandy patches (DS)** | 0.54 | 0.22 | 2.45 | 0.02 |
| **Sperm swimming velocity (10sec after dilution)** | **Density of sandy patches (DS)** | 1.54 | 0.62 | 2.47 | 0.02 |
| ***SPP_1* expressions** | **Floating male (EF)** | -0.92 | 0.89 | -10.2 | 0.3 |
|  | **Courtship success rate (CSR)** | -62.3 | 13.9 | -4.5 | 0.0002 |
|  | **Courtship success rate (CSR) : Floating male (ERFM)** | 23.1 | 4.69 | 4.92 | 0.0007 |
|  | **Courtship success rate (CSR) : Density of sandy patches (DS)** | 9.25 | 3.26 | 2.89 | 0.009 |
|  | **Courtship success rate (CSR) : Floating male (ERFM) : Density of sandy patches (DS)** | -2.64 | 1.1 | -2.37 | 0.02 |
| **GSI** | **Courtship success rate (CSR)** | 0.99 | 0.38 | 2.55 | 0.02 |
|  | **Courtship success rate (CSR) : Floating male (ERFM)** | -0.29 | 0.12 | -2.33 | 0.027 |
|  | **Floating male (ERFM) : Density of sandy patches (DS)** | 0.01 | 0.004 | 2.46 | 0.021 |

**Fig. 3:**
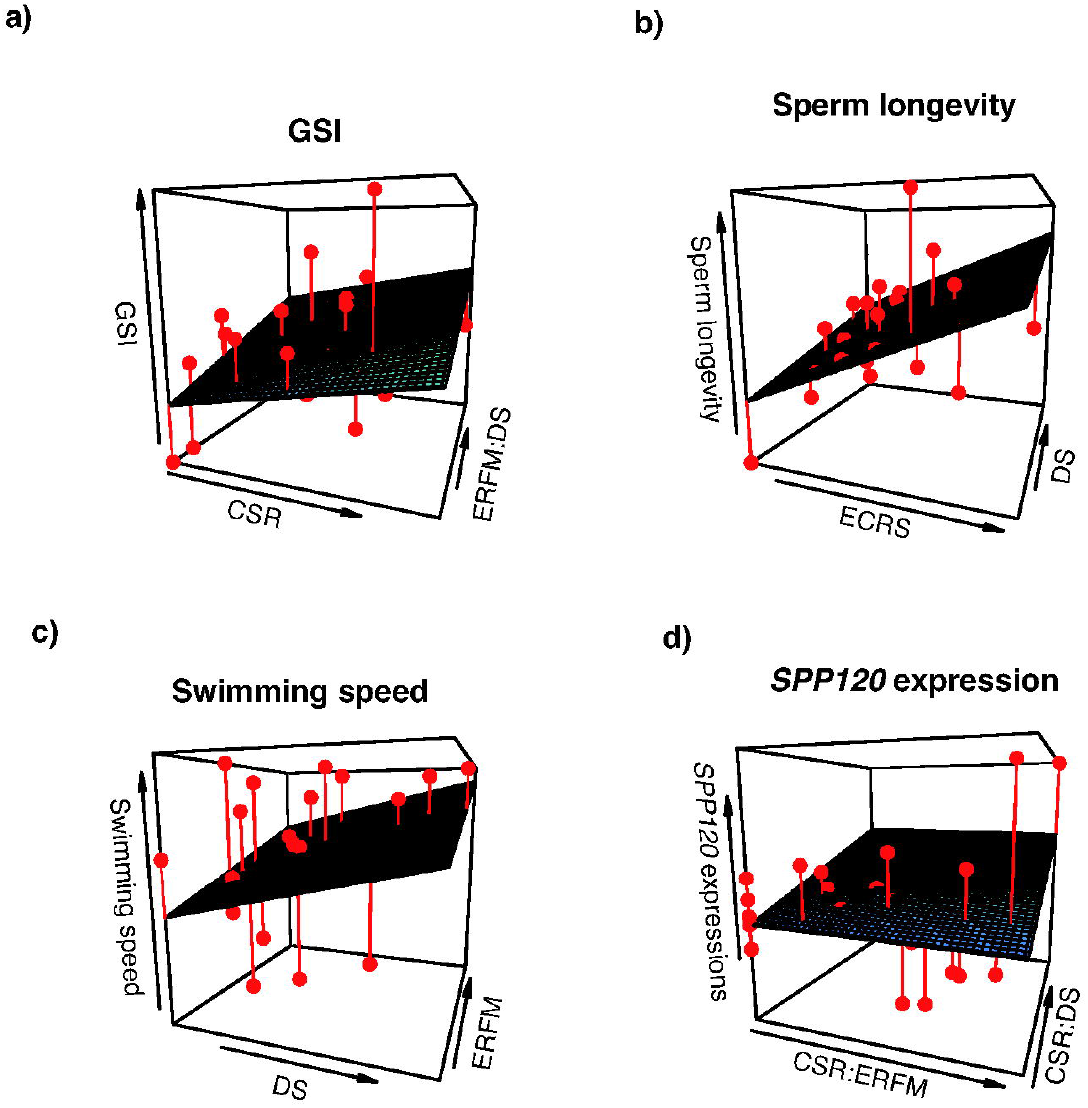
Correlations of sperm quality and behavioral/environmental factors. a) GSI, courtship success rates (CSR), and Encounter rate of floating male (ERFM) and density of sandy patches (DS). b) Sperm longevity, the density of sandy patches (DS), and Encounter rate of floating male (ERFM), c) Sperm swimming speed, the density of sandy patches (DS), and Sperm longevity, d)expression of seminal plasma protein 120 (SPP120), courtship success rates (CSR): Encounter rate of floating male (ERFM), and density of sandy patches (DS): Encounter rate of floating male (ERFM).

Sperm longevity ranged from 133-467 s (Supplementary Fig. 4a) and was positively correlated with DS and ERFM (Table 1, Fig. 3b). Sperm swimming speed varied between 36-61 *μ*m/s (Supplementary Fig. 4b) and was positively correlated with DS and ERFM (Table 1 and Fig. 3c).

The expression of *SPP120_1* was positively correlated with CSR:DS and CSR:ERFM (Table 1, Fig. 3c) but negatively correlated with CSR (Table 1). The expression of *SPP120* pseudogenes was also associated with ERFM and ERFM:DS (Supplementary Table 2). GSI was positively correlated with CSR and ERFM:DS, but weakly negatively correlated with ERFM:CSR (Table 1). Moreover, sperm longevity and swimming speed were not associated (t = 1.18, df = 28, P > 0.05; Supplementary Fig. 5).

## Discussion

From our results, the male can assess the sperm competition level and the sperm quality changes according to the level. For example, the density of the spawning sites and the degree of parasitic spawning raise the sperm competition level. As reported, the involvement of sperm competition changes sperm quality and SPP120 expression. Then, the changes in sperm quality and SPP120 amount can lead to higher fertilization success. However, fertilization preference by females as a representative of the cryptic female choice remains to be understood. Moreover, unpredictable egg release by females was not associated with environmental or behavioral parameters, sperm quality, or SPP amount.

Sperm longevity and swimming speed of *O. ventralis* changed due to phenotypic plasticity and were not correlated with male attractiveness. Indeed, sperm longevity and swimming speed of *O. ventralis* were not associated with CSR, representing male attractiveness (Table 1). For example, *O. ventralis* females prefer to mate with males possessing extended long pelvic fins, which may be genetically determined. However, sperm phenotypic plasticity may not be based on genetics. For example, sperm swimming speed depends on the beat frequency of flagellum associated with intracellular ATP concentration (Chen et al., 2015), and ATP concentration is maintained by many cascades from creatine kinase and mitochondrion (Christen et al., 1983; Tombes and Shapiro, 1985). Furthermore, enzyme activity-dependent ATP concentration in the flagellum is associated with motility (Burness et al., 2004; Perchec et al., 1995; Vignini et al., 2009). The sperm phenotypic plasticity might have been related to differences in ATP amount and enzymatic activity, but a more detailed study is required in this species (see more detail below).

Sperm swimming speed and longevity were not traded in the territorial *O. vetralis* males. In shell-brooding cichlids, *Lamprologus callipterus* (Taborsky et al., 2018), and sea urchins (Levitan, 2000), sperm longevity and swimming velocity change according to sperm competition level and alternative reproductive tactics. For example, in *L. callipterus*, sneaking males have longer-lived sperm, but territorial males have faster-swimming sperm. The differences in sperm motility are reflected in their spawning pattern; territorial males release sperm just after egg release, but sneaking males release sperm before egg release. The energy metabolism fuels flagellar motility leading to different motility patterns according to the alternative reproductive tactics (Hirohashi et al., 2016). The sperm of territorial *O. ventralis* males showed no trade-off between sperm swimming speed and longevity. In squid sperm, the amount of glycogen and environmental conditions for fertilization define their functional differences, while *L. callipterus* dwarf is genetically determined (Wirtz Ocana et al., 2014). However, as discussed above, sperm motility-determinant genetic traits are not linked to dwarf or standard determinant(s).

Sperm quality changes as an outcome of sperm competition, but these changes may be due to a specific mechanism(s) among taxa or their fertilization tactics. We did not observe the sperm motility of the sneaking male during spawning, and hence, further studies are required to elucidate the mechanism(s) involved in generating sperm quality plasticity.

The expression of *SPP120* changes in response to various aspects, and thus, its expression can be influenced by many factors. For example, in our study, SPP120 expression was positively correlated with DS and CSR, which could provoke sperm competition due to multiple sperm shopping. This result is consistent with a previous study, which reported that in mice, the amount of SPP changes according to sperm competition (Ramm et al., 2015).

*SPP120* expression changes according to complex traits associated with sperm competition. For example, male attractiveness and CSR were negatively correlated with *SPP120* expression, but *SPP120* expression increased according to the extent of parasitic males and DS (Table 1). In other words, attractive males (high CSR) do not need to increase *SPP120* expression. Still, their expressions might rise according to the risk of sperm competition caused by parasitic spawning and multiple spawning by females. In Chinook salmon, sperm swimming speed increases in the presence of seminal fluid (Bartlett et al., 2017). Most salmonid fish sperm are motile for a very short time (<60 s) (Turner and Montgomerie, 2002), and their synchronous spawning, often in the presence of sneaking males trying to fertilize, requires faster sperm to achieve fertilization. Similar to that in salmonid fish, in grass goby, the sperm velocity of the sneaker is higher than that of territorial males (Locatello et al., 2013). In *O. ventralis* spawning, the interval between sperm shopping and egg release is unknown for males (Fig. 2b, Supplementary Information 2). In the *O. ventralis*, unlike in the salmonid and goby, SPP120 expression level could be associated with sperm competition level and their spawning manner to raise fertilization success.

Capricious egg deposition in females may not influence sperm quality or *SPP120* expression. In our study, sperm longevity, swimming speed, and *SPP120* expression were not significantly different between males mated with or without egg-released females (Supplementary Fig. 3). On the other hand, the timing of egg release and the number of sperm is predicted to be associated mainly with fertilization success. However, the clutch number of sired males does not correspond with egg deposition in the sandy patches of males because of the cryptic female choice (Haesler et al., 2011). As oral fertilization differs from internal fertilization, the female choice might work through fertilization with sperm from a preferred male. This molecular mechanism (s) can explain fertilization choice for sperm and can confer genetic diversity to the offspring.

In conclusion, sperm competition arises in response to several aspects, and sperm quality and *SPP120* expression are altered due to sperm competition in the mouthbrooding cichlids *O. ventralis*. Their unique fertilization mode (external but enclosed space) is associated with sperm competition. In addition, female behavior (sperm shopping and capricious egg release) and environmental factors, such as DS, influence the sperm competition level. The complex spawning sequences and related factors collectively influence the sperm competition level, thereby changing sperm quality and SPP amount elaborately. In this study, we could not measure the protein amount of SPP120 in the semen and the degree of glycosylation of SPP120. Moreover, it is still obscure how the quality of sperm changes during spermatogenesis. Further research is required to elucidate these aspects to understand the relationship between sperm competition and the plasticity of sperm quality at the molecular level.

## Author contributions

MM conceived the study, conducted field observation, did experiments, analyzed the data, and wrote the paper, MK, SA supported funding, TI and SS supported statistical analyses.

## Data availability statement

Data from this study was uploaded as supplementary data named “Supplemenatry_data_data_set_for_analyses”.

## Figure legends

**Supplementary Figure 1. Schema of the density of sandy patches**. Locations of sandy patches were measured, and we used distances from the patches (dn) to calculate the density of sandy patches (DS).

**Supplementary Figure 2. Maximum likelihood phylogenetic tree and motifs of SPP120 among Tanganyikan cichlids including *Ophthalomotilapia ventralis***. Among *SPP120* copies in the several Tanganyikan cichlids, many copies are found and isolated (Morita et al., 2018). Four copies are found in the examined species *Ophthalmotilapia ventralis*, and two copies are functional.

**Supplementary Figure 3. Comparison among gametic traits and GSI between males who experienced the female egg releases and did not**. We compared a) *SPP120* expressions, b) sperm swimming speed, c) GSI, and d) sperm longevity. We did a Wilcoxon rank sum test to detect differences between those parameters in the males who experienced spawning and did not. There were no differences in every comparison.

**Supplementary Figure 4. Variations of sperm longevity and swimming velocity in the examined individuals**. The variations in sperm longevity and swimming velocity (10 seconds after sperm dilutions) were indicated.

**Supplementary Figure 5. Relationship between sperm swimming speed and sperm longevity**. Sperm swimming speed after 10 seconds dilution of lake water and sperm longevity was shown.

